# 3D light sheet fluorescence microscopy of lungs to dissect local host immune - *Aspergillus fumigatus* interactions

**DOI:** 10.1101/661157

**Authors:** Jorge Amich, Zeinab Mokhtari, Marlene Strobel, Elena Vialetto, Natarajaswamy Kalleda, Katja J. Jarick, Christian Brede, Ana-Laura Jordán-Garrote, Sina Thusek, Katharina Schmiedgen, Berkan Arslan, Jürgen Pinnecker, Christopher R. Thornton, Matthias Gunzer, Sven Krappmann, Hermann Einsele, Katrin G. Heinze, Andreas Beilhack

## Abstract

*Aspergillus fumigatus* is an opportunistic fungal pathogen that can cause life-threatening invasive lung infections in immunodeficient patients. The cellular and molecular processes of infection during onset, establishment and progression are highly complex and depend on both fungal attributes and the immune status of the host. Therefore, preclinical animal models are paramount to investigate and gain better insight into the infection process. Yet, despite their extensive use, commonly employed murine models of invasive pulmonary aspergillosis are not well understood due to analytical limitations. Here we present quantitative light sheet fluorescence microscopy (LSFM) to describe fungal growth and the local immune response in whole lungs at cellular resolution within its anatomical context. We analyzed three very common murine models of pulmonary aspergillosis based on immunosuppression with corticosteroids, chemotherapy-induced leukopenia or myeloablative irradiation. LSFM uncovered distinct architectures of fungal growth and degrees of tissue invasion in each model. Furthermore, LSFM revealed the spatial distribution, interaction and activation of two key immune cell populations in antifungal defense: alveolar macrophages and polymorphonuclear neutrophils. Interestingly, the patterns of fungal growth correlated with the detected effects of the immunosuppressive regimens on the local immune cell populations. Moreover, LSFM demonstrates that the commonly used intranasal route of spore administration did not result in the desired intra-alveolar deposition, as more than 60% of fungal growth occurred outside of the alveolar space. Hence, LSFM allows for more rigorous characterization of murine models of invasive pulmonary aspergillosis and pinpointing their strengths and limitations.

**IMPORTANCE:** The use of animal models of infection is essential to advance our understanding of complex host-pathogen interactions that take place during *Aspergillus fumigatus* lung infections. As in the case of humans, mice need to be immunosuppressed to become susceptible to invasive pulmonary aspergillosis, the most serious infection caused by *A. fumigatus*. There are several immunosuppressive regimens that are routinely used to investigate fungal growth and/or immune responses in murine models of invasive pulmonary aspergillosis (IPA). However, the precise consequences that each immunosuppressive model has on the local immune populations and for fungal growth are not completely understood. Here we employed light sheet fluorescence microscopy (LSFM) to analyze whole lungs at cellular resolution, to pin down the scenario commonly used IPA models. Our results will be valuable to optimize and refine animal models to maximize their use in future research.

**VISUAL ABSTRACT:** Quantitative light sheet fluorescence microscopy to dissect local host-pathogen interactions in the lung after *A. fumigatus* airway infection.

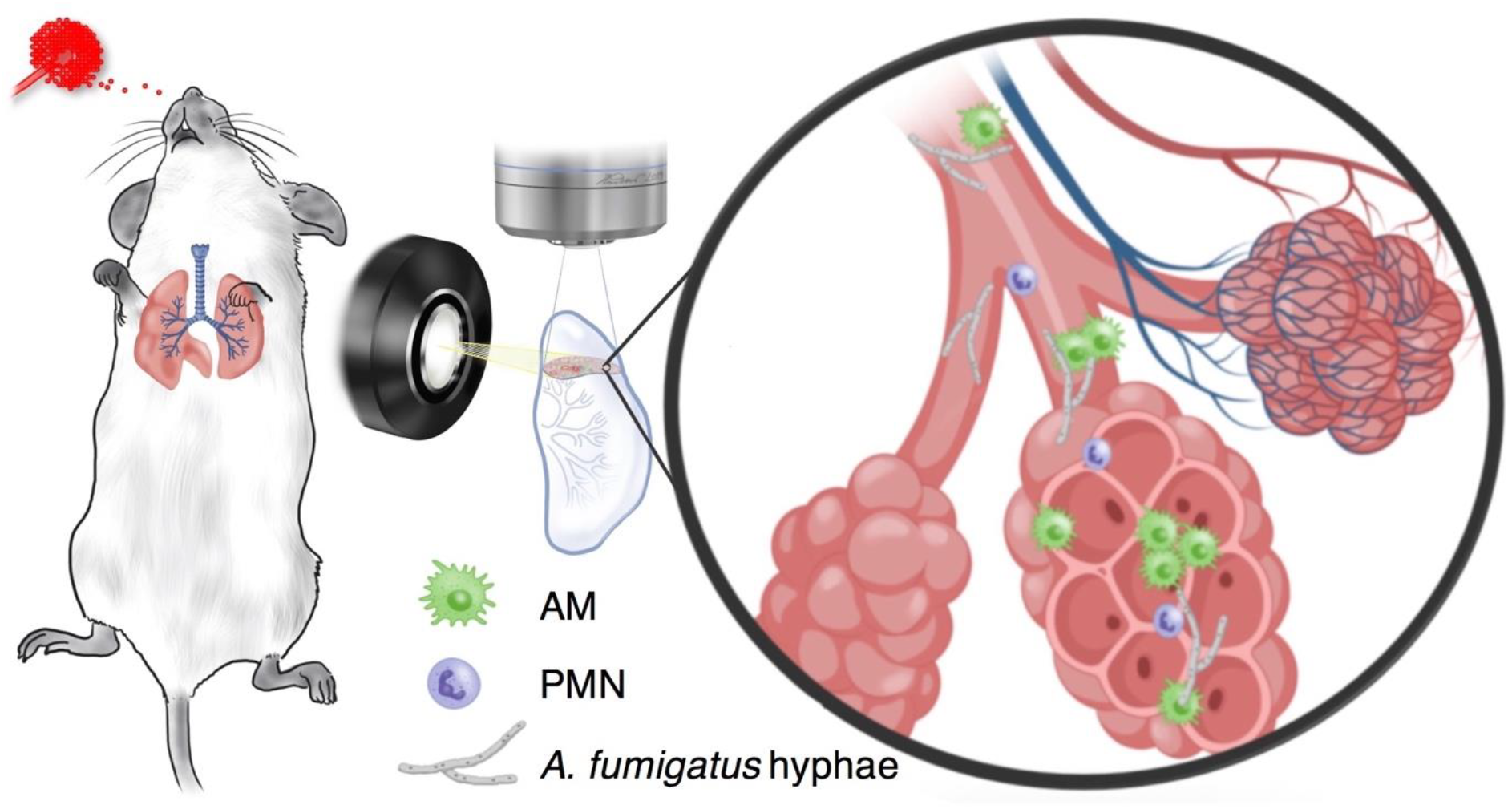

## INTRODUCTION

*Aspergillus fumigatus* is a filamentous, spore-producing, saprotrophic fungus abundant in the natural environment. Airborne *A. fumigatus* spores, termed conidia, can easily penetrate the human respiratory tract and reach the lung alveoli due to their small size (2-3 µm). This rarely has consequences in immunocompetent individuals as innate immune defense mechanisms can very efficiently eliminate conidia. However, *A. fumigatus* can colonize and invade human lung tissues of susceptible hosts with an impaired immune system, causing a spectrum of diseases collectively named aspergillosis. On a world-wide basis, *A. fumigatus* is the most prominent fungal pathogen of the human lung. In Europe, the all-cause burden of *Aspergillus*-related lung disease exceeds 2 million cases annually, including up to 50,000 potentially fatal cases of invasive aspergillosis (IA) [1].

Fungal attributes that enable *A. fumigatus* to thrive in the tissues (metabolic versatility, resistance to stress, etc.) as well as the host immune status that triggers susceptibility (immunosuppression, underlying diseases, etc.) influence the infection process. *In vitro* experimentation has greatly contributed to our understanding of many details in immune–*A. fumigatus* interactions. However, *in vitro* assays cannot achieve a comprehensive picture of the dynamic and complex host pathogen interactions, which justifies studying this multi-layered interplay in *in vivo* models of infection. Particularly, mouse infection models have provided important insights into host fungal *in vivo* interactions and the pathophysiology of the infection process [2]. Yet, precise information about the spatiotemporal evolution of the local host-pathogen interactions has been limited in these animal models. Classic and fluorescence microscopy technologies allow visualization of the interaction of host and fungal cells in the infected organ; however, as the tissue needs to be sectioned into µm-thin slices, the analysis can be only performed on small areas. As a result, analyses of histologic specimens suffer from a sampling bias, the overall three-dimensional anatomical context can be lost, the information obtained is fragmented and the value of data quantification is limited. Non-invasive bioluminescence imaging (BLI) has allowed spatio-temporal tracking of pathogen growth or immune cell recruitment in living animals [3, 4]. However, to date BLI does not provide the optical resolution needed to investigate interactions on a single cell level, consequently often requires complementary approaches for more refined analyses. One technique commonly used is flow cytometry, which permits quantification host cell populations in a high throughput manner [5]. Despite this, analysis of cell populations with flow cytometry requires organs to be processed into single-cell suspensions. Disruption of solid organs and subsequent cell isolation may affect the accurate representation of certain cell subpopulations, often resulting in deviations of extracted cell types, which necessitates high numbers of research animals to reach significant results. Furthermore, flow cytometry is not suitable for the analysis of filamentous fungi and is completely blind for the anatomical context.

A common feature of all *A. fumigatus* infections is persistence and proliferation of inhaled spores in the human respiratory tract. Therefore, to maximize the value of preclinical *in vivo* animal models for improving our understanding of host-pathogen interactions, it is crucial to study the structural organization of fungal spread and the ensuing immune response in the context of the complex spatial microarchitecture of the lung.

Recent advances in light sheet fluorescence microscopy (LSFM) for deep tissue analyses have enormously extended the tool box for studying biological processes [6, 7]. Previously, we employed LSFM to interrogate distributions and interactions in inflammation, cancer and hematopoiesis in whole organs at cellular resolution [8–11]. In the present study, we applied LSFM to investigate the complex spatial relationship underlying *A. fumigatus*-host interactions in lungs of mice under different immunosuppressive regimens. We quantified and visualized fungal growth in the three-dimensional anatomical context of whole lungs. Consequently, we defined the immune response in its anatomical context and quantified host-pathogen interactions within the intact 3D tissue microenvironment of the lung. Following the fundamental tenet that structure and function are inextricably linked, LSFM has proved to be an excellent technique to decipher local host-pathogen interactions in the defined structural context of whole organs at cellular resolution.

## RESULTS

### Light Sheet Fluorescent Microscopy (LSFM) visualization of whole lungs at cellular resolution

To allow for deep-tissue microscopy, we have successfully adapted a previously published protocol for the clearing of whole lungs (Fig. 1A, B) [6, 7]. Employing a custom-built LSFM setup (Fig. 1C), we imaged entire lung lobes with cellular resolution. Intrinsic autofluorescence, detected in the range of 500 - 550 nm, provided anatomical details of the lung airways, revealing the structure of bronchi and bronchioles (Fig. 1D, blue). To detect specific immune cell populations, we employed combinations of fluorescently labelled antibodies. For instance, to visualize CD11c^+^SiglecF^+^ alveolar macrophages (AMs) (Fig. 1E) we detected the combined signal from cells stained with anti-CD11c antibodies conjugated to Alexa Fluor 532 and anti-SiglecF antibodies conjugated to DyLight 755. Additionally, we imaged blood vessels with an anti-CD102 antibody conjugated to Alexa Fluor 488 (A488) to visualize AMs and neutrophil granulocytes (PMNs, CD11b^+^Ly6G^+^) in the context of the pulmonary vasculature. Optical scanning with LSFM allowed to interrogate different depths of the lung tissue (Fig. 1F-H, Supplementary movie 1), and to subsequently rendered series of two dimensional images into a 3D movie (Supplementary movie 2). To detect fungal growth within the lung we utilized the *Aspergillus* specific monoclonal antibody JF5 conjugated with DyLight 650 dye (Fig. 1I-J), which specifically targets an *Aspergillus* mannoprotein antigen secreted during active growth [12]. *A. fumigatus* infection caused disruption of the lung tissue and the appearance of dead cells, which manifested as an increase of tissue autofluorescence and blurring of the observed granular details of well-defined epithelial barriers in healthy mice.

**Figure 1.**
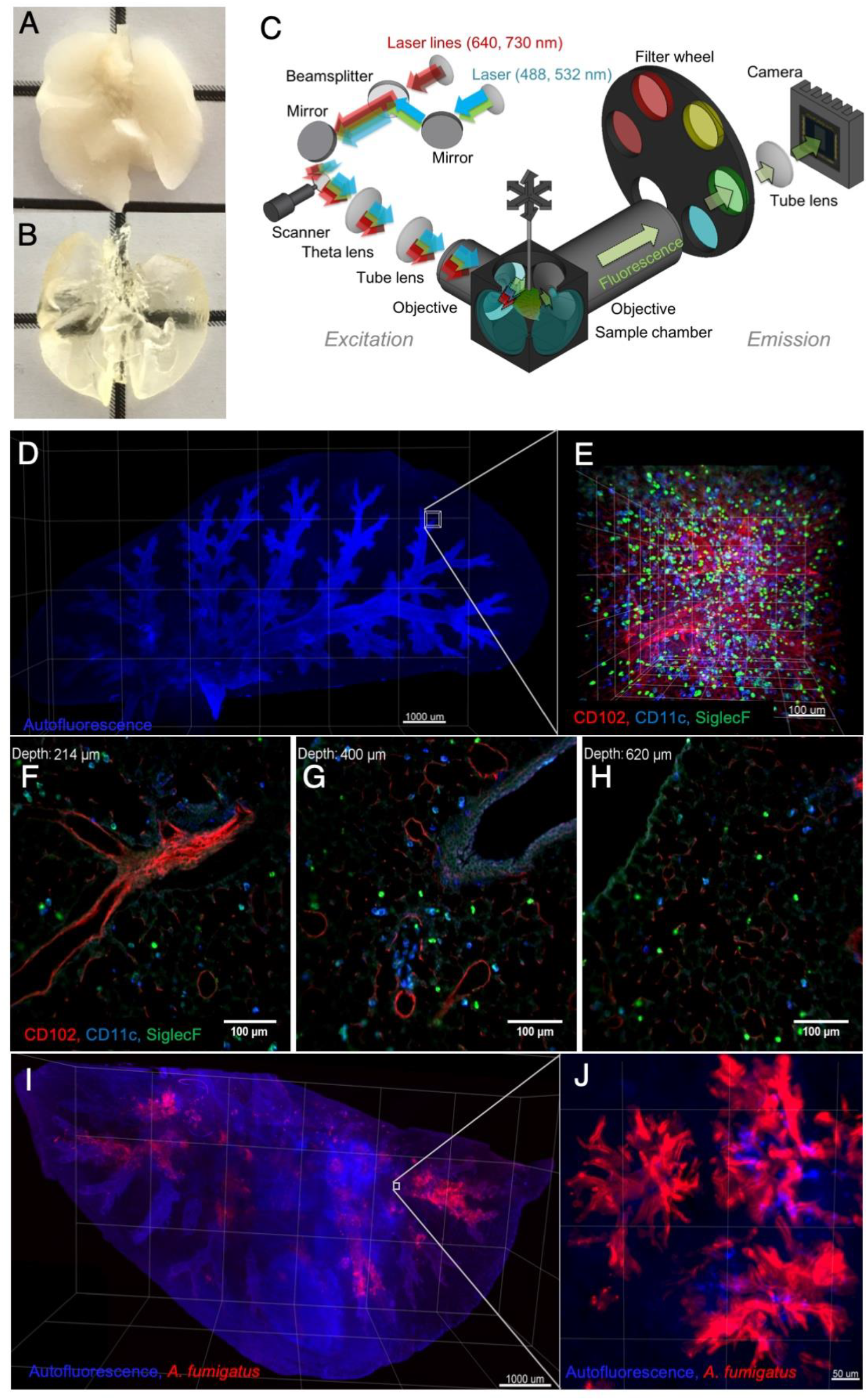
Light sheet fluorescence microscopy (LSFM) maps three-dimensional structural microarchitecture of intact lungs with cellular resolution. **(A)** Whole lungs explanted from a perfused mouse before **(B)** and after clearing. **(C)** Optical setup of LSFM. **(D)** Autofluorescence signal of a murine lung lobe acquired with LSFM. Several fields of view were acquired and stitched together (5X objective, scale bar = 1 mm). **(E)** Zoomed LSFM image depicts immune cell subpopulations at cellular resolution (20X objective, scale bar = 100 µm). Alveolar macrophages (SiglecF^+^ CD11c^+^, light blue) are evenly distributed in the anatomical microenvironment of the lung. CD102^+^ blood vessels are depicted in red. **(F, G, H)** Representative two-dimensional optical sections of the lung of an imaging stack at different penetration depths. **(I)** Lung lobe of a neutropenic mouse infected with 2 × 10^5^ *A. fumigatus* conidia (5x objective, scale bar = 1mm. (**J**) Foci of fungal growth (red) can be observed throughout the tissue as intense cloudy signals at the terminal bronchi (not detected with isotype control antibody). (20X objective, scale bar = 50 µm).

### LSFM reveals the architecture of fungal growth under different immunosuppressive regimens

Murine models of invasive pulmonary aspergillosis (IPA) are the gold standard to investigate the pathogenic potential of different *A. fumigatus* isolates or mutant strains *in vivo* and to evaluate the efficacy of antifungal or immunomodulatory treatments. In mice, as in humans, immunosuppression renders the host susceptible to IPA. To model disease, various immunosuppressive regimens are established, and their use is strongly based on the exact research question (for a review see [13]). The two most prominent immunosuppression models render the mice either leukopenic (usually with alkylating agents), which aims to mimic profoundly neutropenic patients (e.g. suffering from leukemia or undergoing hematopoietic stem cell transplantation) or immunomodulated (usually injecting steroids), which aims to mimic patients treated with immunomodulatory drugs (e.g. after solid-organ transplantation or suffering from graft-versus-host disease (see reference [2] and references within). Despite a basic knowledge of the action of the immunosuppressive treatments, there are still many gaps in our understanding of their effect on fungal growth such as the exact spatiotemporal orchestration of the humoral and cellular interplay recognizing and controlling inhaled fungal spores at physiologic and pathologic conditions. Consequently, we studied three commonly used immunosuppressive regimens (Fig. 2A) to image and quantify the structural organization of fungal growth within intact lungs of *A. fumigatus* infected mice (Fig. 2). Cortisone (C) is known to impair the action of the immune system without depleting AMs or PMNs [14–19]. Notably, under these conditions LSFM revealed only few constrained foci of fungal growth, which covered a small area of the lung, and the fungus could not form proper hyphal filaments (Fig. 2B top). In the most commonly used leukopenic model (CC), mice are treated with the alkylating-agent cyclophosphamide that depletes proliferative immune cells and two doses of cortisone, impairing the function of post-mitotic resident cells [20–28]. Under these conditions, *Aspergillus* invaded the tissue extensively and formed many dense foci of fungal growth that produced long hyphae (Fig. 2B middle). Myeloablative irradiation (Irr) is used as a conditioning treatment for hematopoietic cell transplantation models [4, 29]. This myeloablative conditioning regimen eliminates the host hematopoietic compartment, yet radio-resistant tissue-resident immune cells, such as yolk-sac derived alveolar macrophages, can persist for extended time periods. After myeloablative irradiation we observed that *A. fumigatus* grew extensively, forming also long filaments, yet the numbers of fungal foci were lower than in CC treatment, resulting in a reduced ratio of fungal invasion. Additionally, the structure of the growing masses differed as they formed more dispersed mycelia.

**Figure 2.**
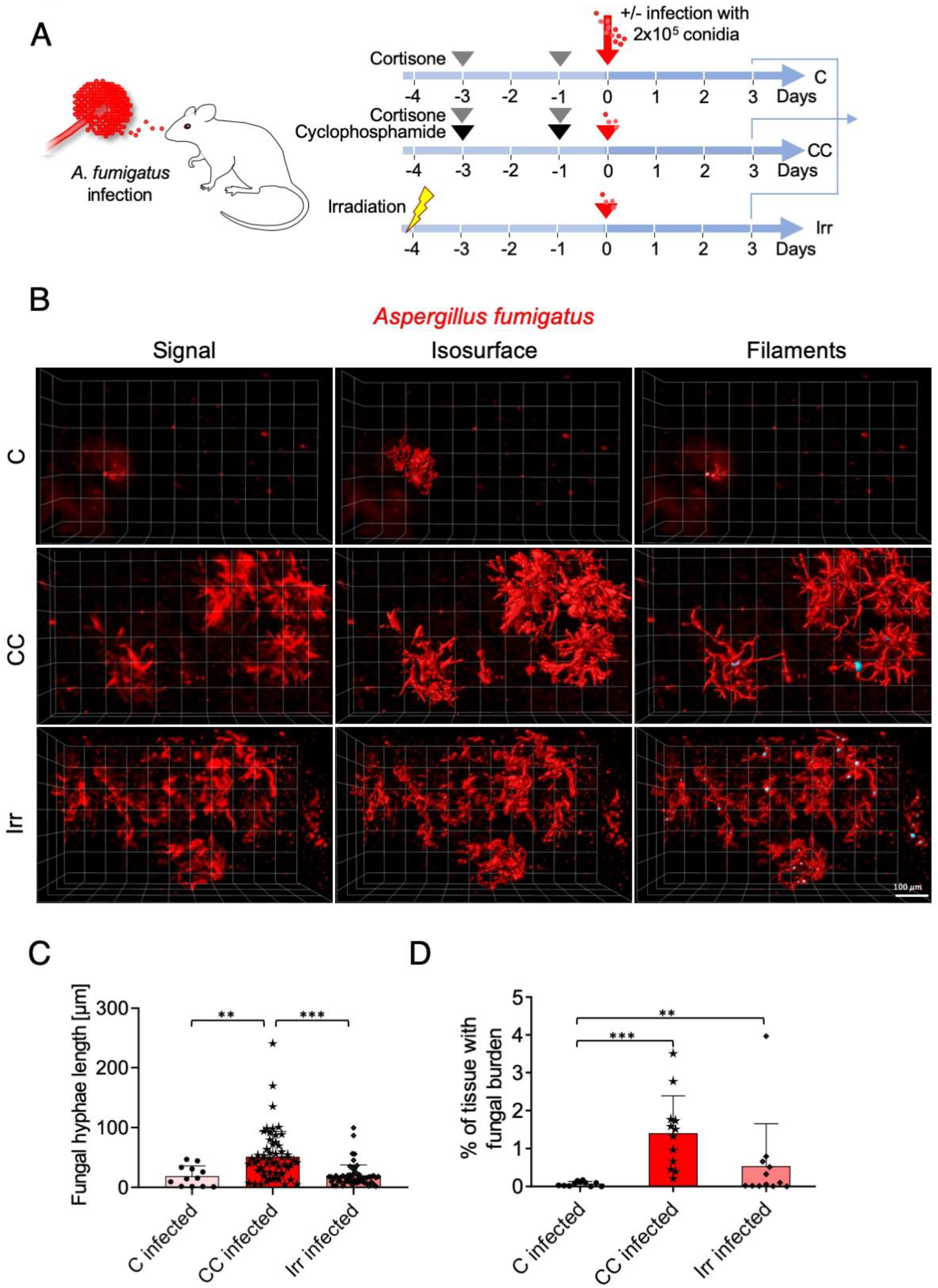
Level of immunosuppression determines local fungal spread. (**A**) Mouse models of immunosuppression employed to study invasive aspergillosis. (**B**) LSFM reveals 3D structures of fungal growth in different models of immunosuppression. Fluorescence signal detected with LSFM (left), calculated isosurfaces of fungal hyphae formation (middle) and filament allocation (right). (20x objective, scale bar = 100 µm (**C**) Length of *A. fumigatus* filaments (hyphae) from the lungs of two mice per condition were analyzed, which triggered measurement of hundreds of filaments per condition. One-Way ANOVA with multiple comparisons was applied. (**D**) Percentage of fungal mass infiltrating lung tissue. Lungs at three randomly selected locations from two mice per condition were analyzed. One-Way ANOVA with multiple comparisons was applied (*** *P < 0.0001*, ** 0.01 < *P <* 0.001).

To this point, LSFM revealed clear differences in fungal growth in the lungs of mice treated with commonly used immunosuppressive regimens and allowed us to map the initiation of IPA in the context of the intact 3D lung environment with high resolution.

### LSFM pinpoints differences in the immune response between immunosuppressive regimens

As LSFM revealed the anatomical information of foci of fungal growth in an unbiased fashion, we explored the specific cellular immune response of AMs and PMNs (Fig. 3A-D) to infection at these hotspots and the impact of common immunosuppressive regimens (Fig. 2A).

**Figure 3.**
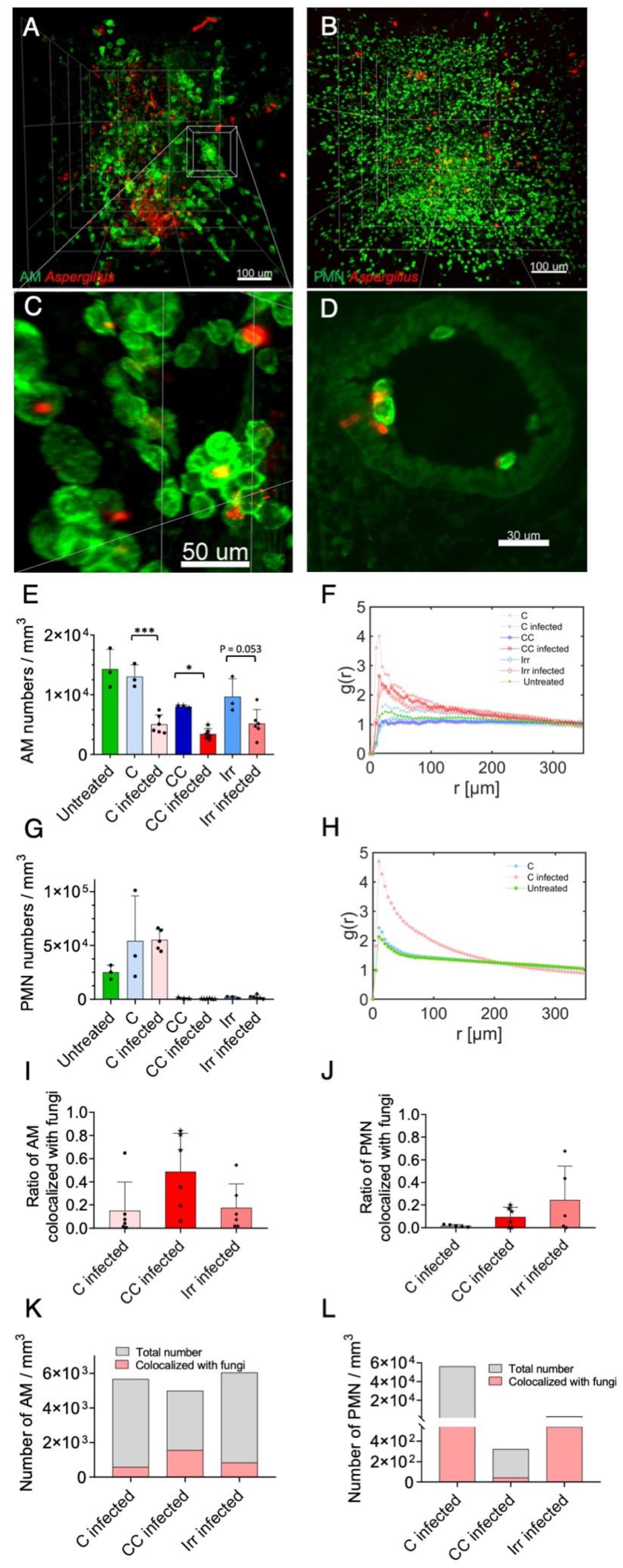
Quantitative image analysis reveals distribution and interaction of immune cells with *A. fumigatus* in 3D lung environment. (**A,B**) Representative 3D images acquired and reconstructed from lungs of infected mice treated with cortisone (C), cyclophosphamide and corticosteroids (CC) or after myeloablative whole body irradiation (Irr). (**A**) AMs (SiglecF^+^ CD11c^+^) depicted in green and *A. fumigatus* (JF5 antibody staining) in red after irradiation and infection. (**B**) PMNs (Ly6G^+^) in infected mouse treated with cortisone are visualized in green. (**C**) Magnification from panel 3A reveals strong AM clustering. (**D**) Fungal spores that had been engulfed by AMs germinate in close vicinity of lung epithelial cells upon infection and irradiation. (**E**) AM numbers quantified with LSFM image analysis in the lungs of mice before and after *A. fumigatus* infection under different immunosuppressive regimens. AM numbers do not decrease after C and Irr treatments and just slightly decline after CC treatment. AM numbers markedly decline upon infection. (**F**) Pair correlation function (g(r)) calculated for AMs using multiple LSFM images from two mice. Correlation reveals that AMs cluster three days after infection, suggesting a recruitment to sites of *A. fumigatus* infection (**G**) Number of PMNs in the lungs of immunosuppressed assessed with quantitative image analysis of LSFM images. The number of PMNs increase after C treatment, which is maintained upon infection. PMNs are virtually eliminated after CC and Irr treatments. (**H**) Pair correlation function for PMNs is calculated only from mice treated with corticosteroids, due to the extremely low numbers in CC and Irr mice. PMNs show a clustered distribution upon infection indicating an active recruitment to the infection sites. (**I-L**) Direct colocalization of AMs and PMNs with *A. fumigatus* indicates that these immune cells ingested fungal material, supporting an active immune response against invasive aspergillosis. One-Way ANOVA Tukey’s multiple comparisons was applied (* *P*=0.04, *** *P<* 0.001).

Corticosteroid treatment did not affect the quantity of AMs in the lungs, but their number significantly decreased upon infection (Fig. 3E), which suggests that many do not survive the challenge with the fungus. Nevertheless, AMs formed clusters upon infection (Fig. 3F), indicating that the corticosteroid treatment did not impair lung resident AMs recognition and interaction with *A. fumigatus*. Corticosteroid treatment significantly increased PMN numbers in the lungs of mice. However, upon infection, their total number did not further increase three days after *A. fumigatus* exposure (Fig. 3G), which suggested that either further PMN recruitment had been impaired or that PMNs died at the same rate as being recruited or relocated to other tissue sites [30]. Notably, LSFM revealed that within lung tissue PMNs strongly clustered upon infection (Fig. 3H), indicating that corticosteroids did not impair the local recruitment and interaction of these cells with the fungus. We then calculated the number of AMs (Fig. 3I-K and PMNs (Fig. 3J-L) that directly co-localized with the fungus as a measure of phagocytic activity. We observed very few AMs and PMNs completely engulfing *A. fumigatus*, suggesting that under corticosteroid treatment these immune cells, although present at the site of infection, were inefficient in eliminating the fungus.

In the leukopenic model, CC treatment slightly decreased AM numbers. *A. fumigatus* infection further reduced AM numbers, suggesting again that AMs died during *A. fumigatus* challenge (Fig. 3E). Similarly, as in the corticosteroid model, CC treatment led to AMs gathering in clusters upon infection (Fig. 3F), indicating that lung resident AMs were not impaired in recognizing *A. fumigatus*. In stark contrast to the corticosteroid regimen, CC treatment virtually eliminated all PMNs in the lungs, and no PMN recruitment after infection indicative of a profound neutropenia. Interestingly, absolute AM numbers (Fig. 3K) that co-localized with the fungus were slightly higher than in the C model, which suggests that AMs killed more *A. fumigatus* conidia in this model. However, PMNs were unable to eliminate resilient *A. fumigatus* spores, which is likely the cause of the huge degree of fungal growth observed in this model (Fig. 2B).

In the irradiation model, infection reduced AM numbers less dramatically compared with the C and CC treatments (Fig. 3E) confirming radiation resistance of AMs even after myeloablative irradiation and infection and indicating many AMs can efficiently eliminate *A. fumigatus* under these conditions. Nevertheless, the number of AMs that had phagocytosed conidia was low (Fig. 3I-K) with some engulfed conidia germinating within AMs (Fig. 3D) indicating that irradiation partially impaired AMs killing potential. In addition, as in the CC model, PMNs were almost entirely depleted from the lungs (Fig. 3G), although the very few that remained seemed to be more efficiently fighting the fungus, as reflected by the higher percentage of PMNs that co-localized with the fungus (Fig. 3J-L), which we propose as the potential reason of the observed reduction in fungal growth (Fig. 2B-D).

Conclusively, LSFM sensitively revealed changes in the total pulmonary lung environment as well as differences in immune cell activity in locally confined foci of *A. fumigatus* infection in subcellular resolution in the lungs of mice treated with various immunosuppressive regimens.

### Majority of A. fumigatus conidia do not reach the lung alveoli in intranasal infection model

One of the factors considered to be important for *A. fumigatus* pathogenic potential is the small size of its conidia (2-3 µm), which permits their penetration through the human respiratory tract to reach the lung alveoli. However, there are morphological differences to the murine respiratory tract, particularly enormous differences in the spatial dimensions, e.g. the distance from the nasal/oral cavities to the tracheobronchial tree or the surface of an alveolus, which is approximately 20-fold smaller in a murine [31] than a human alveolus (estimated mean alveolar volume measures of only < 5.95 × 10^4^ μm^3^ in mice [32] compared to 4.2 × 10^6^ μm^3^ in humans [33]). Therefore, we employed LSFM to reveal spatial information of *A. fumigatus* distribution in the context of the pulmonary microanatomy to address whether the localization of the infective conidia could be affected by anatomical size constrains. To this end, we stained lung epithelial cells with an anti-podoplanin antibody visualizing the epithelial surface to measure the distance of fungal burden to bronchioles by excluding surfaces for the alveoli with diameters less than 50 µm, which is a typical diameter of an alveolus in mice [31, 34] (Fig. 4A). We observed that more than 40% of fungal burden colocalized with bronchioles (first bin of histogram in Fig. 4C) and over 20% located in close vicinity with bronchioles. Only few (~20%) *A. fumigatus* infection foci positioned at sufficient distance (>100 µm) to be considered inside alveoli (Fig. 4C). This proves that around 80% of the *A. fumigatus* conidia did not reach deep into murine lung alveoli upon inoculation, which implies that the murine model of pulmonary aspergillosis may mimic the human infection to a lesser degree than previously anticipated [35]. We further quantified the ratio of AMs (Fig. 4D) and PMNs (Fig. 4F) that located in bronchioles, rather than in the alveoli or other parts of the tissue. Remarkably, we found that a small fraction of AMs localized in the bronchioles in healthy, untreated mice (Fig. 4E), an observation that, to our knowledge, had not been reported until now. Even more interestingly, we found that the ratio of AMs within bronchioles increased upon infection (Fig. 4D), which suggests that AMs were actively attracted out of the alveoli towards the foci of infection in bronchioles. In contrast, under steady-state conditions there were virtually no PMNs in the bronchioles and they were not attracted in C treated mice (Fig. 4F). We documented an increase in CC and Irr mice, but we believe this may rather reflect an artefact caused by the extremely low PMN numbers present in the lungs of these mice.

**Figure 4.**
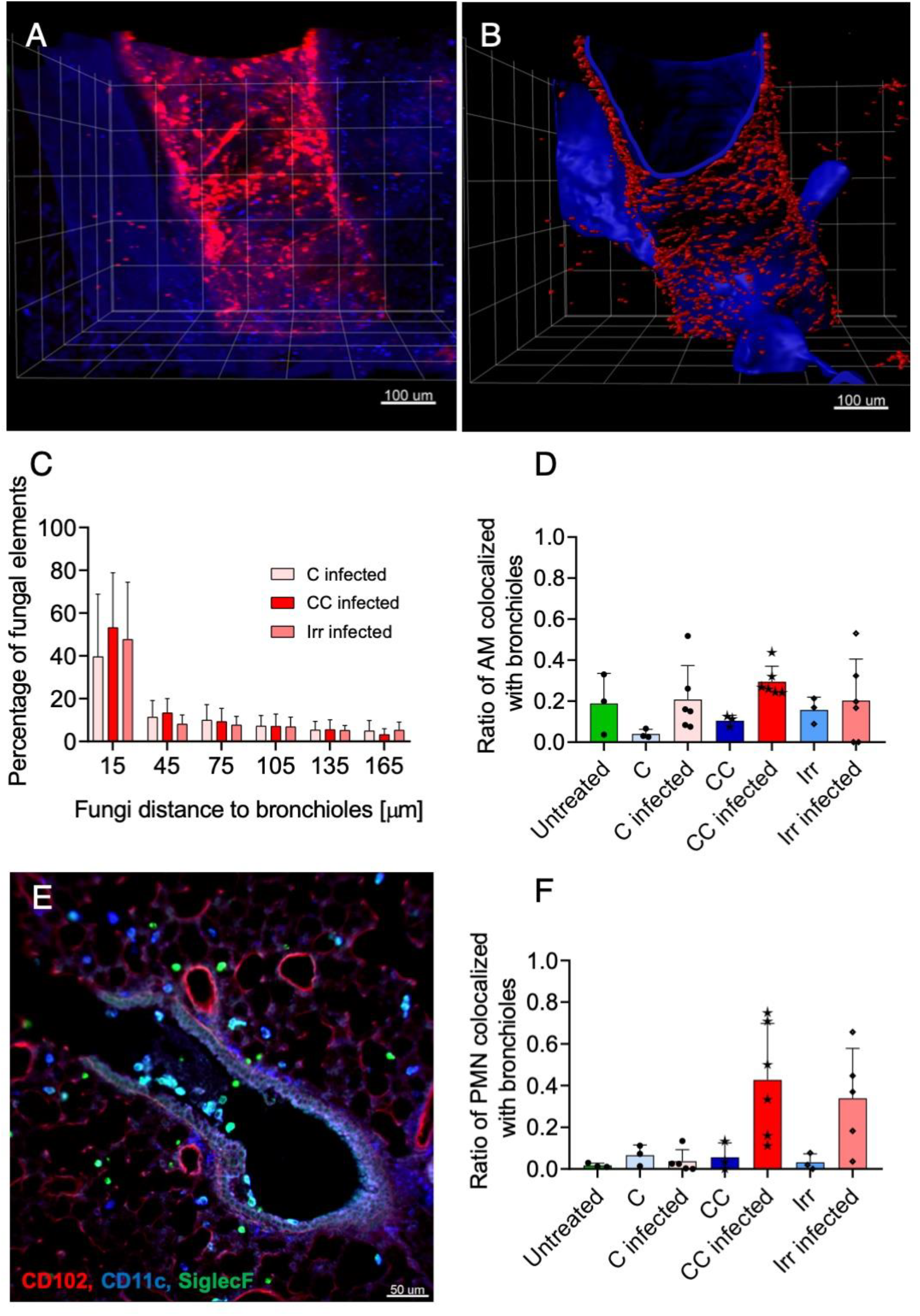
Interaction analysis of fungal burden and bronchioles reveals heterogeneous distribution of fungal spores in murine lungs. (**A,B**) 3D representation of *A. fumigatus* distribution and lung morphology with images acquired and reconstructed from infected lungs **(A).** Fungi and bronchioles isosurfaces **(B)** were reconstructed from anti-*A. fumigatus* JF5 antibody signals and anti-podoplanin staining. Isosurfaces were used to compute distance between fungi and bronchioles of lung. (**C**) Distance distribution of fungi and bronchioles are presented as a histogram for different treatments. In all treatment groups a high percentage of spores (~60%) are in close vicinity with bronchioles, indicating that a vast majority of the infective conidia do not reach the alveoli. Colocalization of AMs (**D-E**) and PMNs (**F**) with bronchioles are calculated and shown as a ratio of total cell numbers. Notably, a low ratio of AMs localized in the bronchioles under steady-state conditions, which increased upon infection for C and CC treated mice. Furthermore, upon infection the ratio of PMNs significantly increased in in CC and Irr treated mice.

In summary, the precise architectural information provided by LSFM uncovered a limitation of the murine model, information that can now be taken into account and tackled by the community. Furthermore, it disclosed a novel location and activity of AMs outside of the alveoli. Hence, LSFM has the potential to open new avenues of investigation in the host-pathogen interplay.

## DISCUSSION

In this study, we employed for the first time LSFM to simultaneously investigate the anatomical distribution of the pathogenic fungus *A. fumigatus* and defined immune cell populations in murine models of pulmonary fungal infection. LSFM proved advantageous in providing quantitative data in the context of spatial anatomical information of large tissue volumes of mouse lungs. Previously, the fungal burden (i.e. the degree of fungal growth) has been measured by colony forming units or by quantitative RT-PCR. Along these lines the number and phenotypic characterization of immune cells in whole lungs has been quantitatively assessed with flow cytometry. However, these methods require homogenization of the lungs and thus destroy the anatomical context of lung tissues. In contrast, LSFM enabled us to quantify fungal burden in the intact tissue environment and to visualize differentiated *A. fumigatus* phenotypes and structures of hyphal growth invading the tissue, thus providing a sensitive measure of the stage of disease development in the context of the tissue architecture. Quantitative microscopy of intact lung lobes provided objective parameters to sensitively measure fungal burden (e.g. expressed as volume % of fungus tissue infiltrates) and differentiate growth stage (e.g. hyphal length). As LSFM can measure even discrete fungal burden within tissues, this method holds great potential to reshape in vivo models to more physiologic infection doses by several orders of magnitude. Beside the scientific merit, this would also contribute to more refined animal experimentation by reducing the disease burden according to the 3R principles (replacement, refinement, reduction for the ethical use of experimental animals). Beyond this, we showed that LSFM can be used concomitantly quantify the number and location of defined immune cell populations. 3D fluorescence microscopy of intact lung lobules provides a clear benefit as it minimizes bias and misinterpretations. Analysis does not suffer from sampling and cell extraction biases and, importantly, provides anatomical data that allows calculating the distribution of the cells in the tissue, their clustering with *A. fumigatus* and their phagocytosis rate upon infection. Until recently, the only alternative to obtain lung anatomical information was to perform histology of thin consecutive lung sections, staining them with certain dyes (for instance PAS or Gomori-Grocott for fungal hyphae or specific antibodies (to detect, for example, specific cell populations [36]). Analyses of consecutive histological specimens are time and labor intensive and, therefore, the volume of tissue that can be investigated within a reasonable time frame is small. Thus, histology sections serve as a reference of fungal germination/hyphal enlargement or presence/absence of immune cells, but provide qualitative rather than quantitative data. Recently, Shevchenko and colleagues utilized confocal microscopy to investigate neutrophil recruitment and location in the conducting airway upon *A. fumigatus* infection [37]. Although this is a significant improvement, this technique is still limited by the small fraction of tissue imaged, which makes any quantification and anatomical information relative. LSFM permits to image whole lung lobes of infected mice and, therefore, to capture the entire host-pathogen interaction. Nevertheless, LSFM has some limitations as well. For example, sample preparation may be difficult, when mice need to be perfused, time-consuming as staining and clearing procedures take several days, and expensive with high amounts of antibodies being required. Consequently, while this technique has enormous potential, it is still in its infancy when it comes to high throughput or exploratory analyses. In addition, the advantage of LSFM in whole-organ imaging is off-set by lower spatial resolution compared to confocal microscopy [12, 38, 39], which enables more refined subcellular analyses [40]. In addition, LSFM of the opaque lung tissue also requires tissue fixation, and thus only allows snapshots of the infection process. In contrast, multiphoton microscopy allows for live imaging of explanted organs [39, 41–43] or even *in vivo* microscopy [44, 45] using a thoracic window [46] even if a smaller field of view. Fortunately, as recently demonstrated [11], LSFM and two-photon microscopy can complement each other with high added value so that both tissue architecture and cell dynamics can be interrogated to investigate the dynamic temporal host-pathogen interactions as well as its large-scale distributions.

In this study, LSFM proved especially advantageous to visualize and quantify fungal growth and the immune response in the lungs of mice treated with various immunosuppressive regimens commonly used to model invasive pulmonary aspergillosis. It has previously been reported that *A. fumigatus* extensively grows and invades the lung tissue in cyclophosphamide and cortisone (CC) treated mice, whilst its growth is restricted in cortisone (C) treated mice [18, 47–49]. However, this had previously only been inferred from classical histology sections, as the actual 3D structure of fungal foci in the whole lungs had not been visualized. Here we show that in corticosteroid treated mice *A. fumigatus* formed condensed foci of growth without hyphal extensions throughout the tissue and the percentage of invaded lung tissue remained very low. In contrast, in CC mice the fungus created a complex 3D hyphal structure that extended throughout the surrounding tissue invading a high percentage of tissue volume (average of 1.5% of the whole lung). Interestingly, although fungal hyphae formed in irradiated mice, these comprised only an intermediate fungal architecture that invaded lung tissue at a lower level than observed in CC mice. Such fungal growth patterns matched with the observed effects of the immunosuppressive drugs on AMs and PMNs [50, 51]. Neither cell population was depleted in mice treated with corticosteroids, and as result fungal growth was constrained. However, AMs’ function appeared to be compromised revealed by AM numbers directly colocalizing with *A. fumigatus* and infection did not further increase the already high PMN numbers. Therefore, the sustained attempt to eliminate persisting fungus likely contributes to the hyper-inflammation characteristics of this model [48, 52, 53]. In the CC model AM numbers were similar to mice receiving only corticosteroids; this demonstrates that cyclophosphamide treatment did not deplete AMs. Upon infection, AMs also clustered with *A. fumigatus*, suggesting that CC treatment even did not affect the recognition and interaction of AMs with *A. fumigatus*. Notably, the ratio of AMs interacting with *A. fumigatus* conidia was higher in CC mice than in C mice. It is tempting to speculate that this might be explained by the lack of a coordinated PMN defense leaving AMs as the only cell type to interact with conidia. Therefore, it seems that AMs as first responders attempt to contain the infection. If AMs cannot eliminate *A. fumigatus*, the host relies on an efficient secondary response by PMNs to control the fungus. If PMNs are completely abrogated, as observed in the scenario of CC mice, massive fungal growth culminates in an invasive aspergillosis. In lethally irradiated mice, *A. fumigatus* grew more extensively than in the corticosteroid model, likely because there were not sufficient PMN numbers to act as second line of defense; however, growth was slightly less than in the CC model, which could be explained by the fact that AMs and the few PMNs present seemed to remain mostly functional to eliminated many conidia [54]. Accordingly, irradiated mice displayed fewer foci of fungal growth than CC treated mice and the more dispersed mycelia indicated that remaining host immune cells contained the growing hyphae to a certain degree.

The significant differences in the dimensions of the murine airways [55] compared to humans affect the distribution of inhaled conidia. LSFM revealed that about 80% of the intranasally administered *A. fumigatus* spores did not reach deep into the lung alveoli. This finding has important implications for the design of murine experiments and the conclusions that can be reached from them. For example, elastase has been disregarded as a virulence factor in *A. fumigatus* [56], but that could be because degrading elastin might be more relevant in the alveoli, where elastin is readily present in the alveolar wall. Besides, the first contact with the host will not be orchestrated exclusively by alveolar epithelial cells, as it is normally assumed in humans [57]. Additionally, *in vivo* murine studies may result in underestimating the role of AMs for human infection [49]. Therefore, other methods that ensure a better distribution of the spores could be more accurate to define the features of IPA [35]. Inhalation models have been already described since the early 90’s [58] and their value has been proven [59]. A model of intratracheal intubation with ventilation has also been described to trigger a better distribution of spores [12, 60]. Hence, we believe that such models should be better characterized to assess whether these should be adopted as the gold standard for pulmonary infection models.

Interestingly, we have observed that a small ratio of CD11c^+^SiglecF^+^ AMs localized outside of the alveoli, in the bronchiolar space, an observation that, to the best of our knowledge, had not been reported before. Upon infection, the ratio of AM in bronchioles increase, suggesting an active recruitment towards foci of fungal infection outside of the alveolar space. We believe this is an important result that may open new research avenues and interesting questions. Are these cells a distinct AM subset? How are these cells recruited to these sites? What are the distances they can overcome? Which signals attract them?

In summary, we have shown that state-of-the-art LSFM is a powerful tool to describe important features of fungal growth and of the local immune response in commonly used murine models of pulmonary aspergillosis. We have demonstrated that LSFM can comprehensively localize all individual foci of fungal growth and permits to investigate the local fungal-pathogen interactions at cellular resolution. This opens the possibility to assay models of pulmonary aspergillosis using lower, more physiological, doses of infective conidia and still gain insight into the immune response. Hence this study not only advances our understanding of the infection process under immunosuppression, but also could form the basis to refine and maximize the utility of animal models of pulmonary infection.

## MATERIAL AND METHODS

### Mice

All experiments were conducted with female 8 to 12-weeks-old BALB/c mice (Charles River, Sulzfeld, Germany). Mice were maintained in individually ventilated (IVC) cages with *ad libitum* access to water and food. All experiments were performed according to the German regulations for animal experimentation and approved by the Regierung von Unterfranken (55.2-2531.01-86-13 and 55.2-2532-2-403) as responsible authority.

### Immunosuppression regimens and A. fumigatus infection

Mice were injected subcutaneously with 112 mg/kg hydrocortisone acetate (Sigma-Aldrich) alone (C group) or together with 150 mg/kg cyclophosphamide (Sigma-Aldrich) intraperitoneally on days –3 and –1 prior to infection (CC group). Mice were myeloablatively irradiated (8 Gy) with an electron linear accelerator (Mevatron Primus, Siemens, Germany) 3 hours before infection (Irr group).

On day 0, mice were intranasally infected with 2 × 10^5^ freshly harvested conidia (suspended in 0.9% NaCl + 0.005% Tween-20) of the *A. fumigatus* clinical wild-type isolate ATCC46645.

### Lung preparation for LSFM

Prior to lung extraction, mice were perfused using an ISMATEC Reglo Analog pump (IDEX Health & Science LLC, Oak Harbor, USA) 2 minutes with PBS and 8 minutes with paraformaldehyde (PFA) 4%. Lungs were further fixed 2 hours in 4% PFA and washed 3×30 minutes with PBS before processing. To visualize *Aspergillus fumigatus*, lungs were blocked overnight in PBS / 2% FCS / 0.1% Triton-X and monoclonal antibody JF5 (DyLight 655) against aspergillus fumigates were added in a 1:100 dilution. To stain immune cell populations in mouse lungs the following antibodies were used: Anti-CD11b (M1/70) (AF488), anti-CD11c (N418) (AF488), anti-Ly6G (1A8) (DyLight 755) anti-SiglecF (E50-2440)(DyLight 755) from Biolegend (Uithoorn, The Netherlands) and eBioscience (Frankfurt, Germany). All antibodies were added in a 1:100 dilution in PBS to the samples and were incubated 2 days at 4 °C under gentle shaking. After staining, lungs were washed 3×30 minutes with PBS, samples were dehydrated in a graded ethanol series (30%, 50%, 70%, 80%, 90% for 1.5 hours each at room temperature, and in 100% overnight at 4 °C). Next day the samples were rinsed for 2 hours in 100% *n*-hexane; afterwards *n*-hexane was replaced stepwise by a clearing solution consisting of 1 part benzyl alcohol in 2 parts benzyl benzoate (Sigma-Aldrich). Air exposure was strictly avoided at this step. Tissue specimens became optically transparent and suitable for the LSFM imaging after incubation in the clearing solution for at least 2 hours at room temperature.

### LSFM setup and data acquisition

The LSFM setup is home-built. For excitation a customized fiber coupled laser combiner (BFI OPTiLAS GmbH, Groebenzell, Germany) was used that provided the required excitation lines of 491, 532, 642, and 730 nm. For laser beam collimation (beam diameter ≅ 3 mm) a Hund objective (A10/0.25 Hund, Wetzlar, Germany) was used. A dichroic beam splitter DCLP 660 (AHF Analysentechnik, Tübingen, Germany) combined both beam paths. A single axis galvanometer scanner (6210H, Cambridge Technologies, Bedfort, MA, USA) in combination with a theta lens (VISIR f. TCS-MR II, Leica, Mannheim, Germany) finally created a virtual light sheet that was additionally pivot-scanned by a two-axis resonant scanner system (EOP-SC-20-20×20-30-120, Laser2000, Wessling, Germany) to minimize shadowing artefacts [61]. The light sheet was projected onto the sample via a tube lens and an objective lens (EC Plan-Neofluar 5x/0.16 M27, Zeiss, Göttingen, Germany). The objective on the detection side (HCX APO L20x/0.95 IMM Leica, Mannheim, Germany) collected the fluorescence perpendicular to the light sheet, and, in combination with an infinity-corrected 1.3x tube lens ∞/240-340 (098.9001.000, Leica, Mannheim, Germany) projected the image into a sCMOS camera Neo 5.5 (2560 x 2160 pixels, 16.6 mm x 14.0 mm sensor size, 6.5 µm pixel size, Andor, Belfast, UK). The fluorescence was spectrally filtered by typical emission filters (AHF Analysentechnik, Tübingen, Germany) according to the used fluorophores as follows: BrightLine HC 525/50 (Autofluorescence), BrightLine HC 580/60 (Alexa Fluor 532), HQ697/58 (Alexa Fluor 647), BrightLine HC 785/62 (Alexa Fluor 750). Filters were part of a motorized filter wheel (MAC 6000 Filter Wheel Emission TV 60 C 1.0x (D) with MAC 6000 Controller, Zeiss, Göttingen, Germany) placed in the collimated light path between detection objective and tube lens. Multicolor stacks were acquired in increments of 2 µm by imaging each plane in all color channels sequentially. Hardware components for image acquisition (laser, camera, lenses) were controlled by IQ 2.9 software (Andor, Belfast UK). Images were saved as tiff-files and analyzed as described below.

### Analysis of LSFM images

IMARIS software v8.1.1 (Bitplane AG, California, USA) was employed to analyze the images obtained with LSFM. When required, a background subtraction in accordance with the diameter of the cell population was applied to eliminate unspecific background signals. Fungal and cellular isosurfaces were segmented using the option “Surface” with a smoothing of 10% of the expected diameter of the object and manual thresholding of the signal. Filaments were calculated from a masked channel from the *A. fumigatus* isosurfaces, using starting points of 50-70 µm and seed points 7-3 µm (depending of the degree of fungal invasion) and manual thresholding of the signal. To measure the distance of fungal burden from bronchioles, we have created isosurfaces from lung epithelial cells (podoplanin) signal and filtered surfaces with diameter less than 50 µm to exclude alveolus structures.

### Cell distribution analysis

The distribution of stained cells in the light sheet microscopy data resembles a spatial point pattern in mouse lungs [62]. To identify the clustering of the cells and characterize the cell distribution patterns, pair correlation function (g(r)) [63, 64] was calculated at distance r as following: g(r) = average number of cells within rings at distance r from an arbitrary cell. The pair correlation function can correctly identify the aggregation length scale and distance between clusters of cells in the lung. In case of having random cell distribution, g(r will be about 1. Cells expulsion can be revealed by g(r)S< 1, and aggregation is realized by g(r) > 1.

### Statistical analysis

Ordinary one-way ANOVA test with Tukey’s multiple comparisons was used for normally distributed data. Non-normal data was compared employing Kruskal-Wallis test. All statistical analysis was performed with GraphPad Prism Version 7.

## ACKNOWLEDGEMENTS

This work was supported by the German Research Council to AB and KGH (DFG TRR124 A3). We thank the members of the Beilhack laboratory, the Heinze laboratory, Drs. A. Brakhage and M. Brock and the members and the TRR124 consortium for vivid discussions.

## AUTHOR CONTRIBUTIONS

JA, ZM and AB designed the study. KGH and JP built and customized the LSFM setup. JA, ZM performed experiments and analyzed the data. JA, ZM, KGH, AB prepared the figures and wrote the paper. MS, EV, NK, KJJ, CB, ALJG, ST, KS, BA, JP supported experiments and data analysis. HE, CRT, MG, and SK provided important reagents and contributed to manuscript preparation.

## SUPPLEMENTARY MOVIES

**Supplementary movie 1:**

**Optical sectioning of an intact lung lobe with multicolor LSFM.**

**Supplementary movie 2:**

**Virtual journey through an intact lung lobe with multicolor LSFM of healthy and A. fumigatus infected mice.**

